# Resistance of *Musa balbisiana* accessions of the Philippines to *banana bunchy top virus*

**DOI:** 10.1101/2022.08.20.504392

**Authors:** Fe M. Dela Cueva, Nicole Angelee M. Perez, Ann Fatima A. Benjamin, Lyka A. Yanos, Lavernee S. Gueco, John E. Thomas

## Abstract

Mitigation of banana bunchy top disease (BBTD) is still a challenge worldwide. BBTD is caused by banana bunchy top virus (BBTV), the most important virus affecting banana. Currently, no cultivar or accession of banana has complete resistance to BBTD. A total of 36 wild *Musa* spp. accessions, including 34 *Musa balbisiana* and two *M*. *acuminata* subsp. errans (‘Agutay’), were screened for resistance against BBTV. In greenhouse tests using viruliferous banana aphids *(Pentalonia nigronervosa*), all *M. balbisiana* accessions remained symptomless and BBTV was not detected in any of these plants by PCR at three- and six-months post inoculation. In contrast, 100% disease incidence was recorded in *M. acuminata* subsp. *errans* and in cv. Lakatan susceptible control plants. The PCR-negative *M*.*balbisiana* plants were then transferred to a field with high BBTV inoculum pressure where they remained symptomless and PCR-negative for up to five years, while all cv. Lakatan developed BBTD. This study confirmed the resistance of wild *M*.*balbisiana* accessions to BBTV. It is therefore expected to provide a resource for conventional and marker assisted breeding.

## Introduction

Banana bunchy top disease (BBTD) continues to spread and affect banana production worldwide. Recent records include the first reports of BBTV from Tanzania (Mpoki et al., 2021; and Uganda (Ocimati et al., 2021). The occurrence of this disease in banana cropping areas planted with susceptible varieties can result in a total yield loss (Qazi, 2015). Once infected with BBTV, plants rarely recover. Banana bunchy top virus (BBTV; genus *Babuvirus*, family *Nanoviridae*) is the causal agent of BBTD and its genome comprises six circular ssDNA components (Harding et al., 1991; Xie & Hu, 1995). The major vector is the banana aphid or *Pentalonia nigronervosa*(Dela Cueva et al., 2015; Thomas, 2008). Long-distance transmission is attributed to the movement of infected planting materials (Thomas et al., 1994).

BBTV occurs widely in Africa, Asia, and the Pacific and is difficult to control. Common strategies against the disease are quarantine, eradication of infected plants, and the use of clean planting material (Thomas, 2019). Widely used control practices in the Philippines is eradication of infected plants using bamboo sticks dipped in herbicide, and the use of clean planting materials (Aguilar et al., 2010; Molina et al., 2009). A number of studies have explored the potential of transgenic resistance to BBTV in bananas (Borth et al., 2011; Shekhawat et al., 2012; Elayabalan et al., 2013; Mware et al 2016) but to date, these advances have not been successfully transferred to a field situation. Biopriming with endophytic and rhizosphere bacteria (Kavino et. al., 2007; Harish et al., 2009; Jebakumar and Selvarajan 2018) has shown some promise for BBTD control when applied prior to infection, resulting in reduced incidence of infection and a delay in symptom expression under greenhouse conditions. Screening and breeding of new *Musa* spp. cultivars have been done in attempts to improve resistance to important banana pests and diseases (Ortiz, 2013), but BBTV still poses a challenge as there are still no known natural sources of resistance against this virus (Shekhawat et al., 2012).

The wild species of banana, *M*. *acuminata* (AAw) and *M*.*balbisiana* (BBw), are the major sources of the genomes of edible bananas cultivated today, contributing “A” genome and “B” genome components, respectively (Valmayor et al., 2002; Venkataramana et al., 2015). Though ultimately susceptible, banana cultivars can differ markedly in the ease of infection with BBTV and with the intensity of symptoms expressed and it has often been observed that banana cultivars with a B genome component tend to display a level of resistance to BBTV (Ngatat et al., 2017; Jose, 1981; Muharam; 1984, Espino et al., 1993; Niyongere et al., 2011; Sachter-Smith, 2015; Hapsari and Masrum; 2012, Latifah et al., 2021). Resistance against important diseases has been sought in genotypes of banana from the wild (Jenny et al., 2002) though none have been conclusively established for BBTV. The Philippines has a diverse collection of species of banana from the wild (Valmayor et al., 2002). Recognizing this potential, this study screened Philippines wild-type banana species, predominantly *M*.*balbisiana*, from the germplasm collection of National Plant Genetic Resources Laboratory, Institute of Plant Breeding (NPGRL, IPB) for resistance against BBTV.

## Materials and methods

### Study Site

The screening trial of wild banana was conducted in a greenhouse (14°9’19”N, 121°15’40”E) and field research site (14°8’60”N, 121°16’20”E) at the Institute of Plant Breeding, College of Agriculture and Food Science, University of the Philippines-Los Baños (IPB, CAFS, UPLB). The study started in November 2015 and the experimental field was selected due to the prevalence of BBTV and banana aphid in the area. The field trial site occupied 7,950 m^2^ with nine experimental plots. Plant spacing was 1m within rows and 2m between. The area had an average rainfall of 1700 mm and annual mean temperature of 23.8°C (minimum) and 31.7°C (maximum) from 2016-2021.

### Preparation of Musa accessions

Thirty-six wild banana accessions of the Philippines were screened in the study. It includes 34 Musa balbisiana accessions and two M. acuminata subsp. errans. The wild banana progenitors were collected from different regions of the Philippines as illustrated in figure 1. The collection site and identification of each accession are specified in table 1. The two *M*. *acuminata* subsp. *errans* and 32 *M*. *balbisiana* accessions were derived from seeds. The remaining two *M*. *balbisiana* are tissue-cultured. Test plants from seeds and tissue culture were transplanted to pots with sterilized soil and coir dust at a ratio of 1:3. Test plants were incubated under greenhouse conditions. Two-month-old seedlings were used in artificial inoculation of BBTV.

**Table 1.** List of wild Musa accessions screened from germplasm collection of NPGRL-IPB

| Plant Source Type | GB/ PHL Number | Collection Number | Local name | Scientific name | Collection Site |
| --- | --- | --- | --- | --- | --- |
| Seeds | GB24350 | 1997-276 | Butuhan | *Musa balbisiana* | Zambales |
| Seeds | GB24364 | 1998-078 | Butuhan | *Musa balbisiana* | Quirino |
| Seeds | GB24366 | 1998-085 | Butuhan | *Musa balbisiana* | Quirino |
| Seeds | GB24376 | 1998-315 | Pik-iw | *Musa balbisiana* | Marinduque |
| Seeds | GB24376 | 1998-315 H2 | Pik-iw | *Musa balbisiana* | Marinduque |
| Seeds | GB24405 | 1998-453 | Butuhan | *Musa balbisiana* | Ilocos Norte |
| Seeds |  | 1998-456 | Balibisiana | *Musa balbisiana* | Ilocos Sur |
| Seeds | GB24421 | 1998-575 | Butuhan | *Musa balbisiana* | Leyte |
| Seeds | GB24442 | 1998-633 | Pik-iw | *Musa balbisiana* | Quezon |
| Seeds | PHL32673 | 2010-047 | Balbisiana | *Musa balbisiana* | Oriental Mindoro |
| Seeds | GB61941 | 2013-100 | Pakol | *Musa balbisiana* | Catanduanes |

| Plant Type | GB/ PHL Number | Collection Number | Local name | Scientific name | Collection Site |
| --- | --- | --- | --- | --- | --- |
| Seeds | GB61941 | 2013-100 H2 | Moko | *Musa balbisiana* | Catanduanes |
| Seeds | GB61942 | 2013-101 | Butuhan | *Musa balbisiana* | Catanduanes |
| Seeds | GB61996 | 2013-155 | Moko-Bulo | *Musa balbisiana* | Camarines Sur |
| Seeds | GB61996 | 2013-155 H2 | Moko | *Musa balbisiana* | Camarines Sur |
| Seeds | GB61997 | 2013-156 | Butuhan | *Musa balbisiana* | Camarines Sur |
| Seeds | GB62000 | 2013-159 | Moko | *Musa balbisiana* | Camarines Norte |
| Seeds | GB66554 | 2016-003 | Pik-ew/ Butuhan | *Musa balbisiana* | Agusan del Norte |
| Seeds | GB66560 | 2016-013 | Balbisiana | *Musa balbisiana* | Oriental Mindoro |
| Seeds | GB66565 | 2016-018 | Balbisiana | *Musa balbisiana* | Agusan del Norte |
| Seeds | GB66565 | 2016-018 | Balbisiana | *Musa balbisiana* | Agusan del Norte |
| Seeds | GB66583 | 2016-048 H1 | Pakol | *Musa balbisiana* | Albay |

| Plant Source Type | GB/ PHL Number | Collection Number | Local name | Scientific name | Collection Site |
| --- | --- | --- | --- | --- | --- |
| Seeds | GB66586 | 2016-051 | Butuhan | *Musa balbisiana* | Camarines Sur |
| Seeds | GB66586 | 2016-051 H1 | Butuhan | *Musa balbisiana* | Camarines Sur |
| Seeds | GB66586 | 2016-051 H2 | Butuhan | *Musa balbisiana* | Camarines Sur |
| Seeds | GB66587 | 2016-052 | Balibisiana | *Musa balbisiana* | Camarines Sur |
| Seeds | GB66588 | 2016-053 | Balibisiana | *Musa balbisiana* | Camarines Sur |
| Seeds | GB66591 | 2016-056 H3 | Balbisiana | *Musa balbisiana* | Camarines Norte |
| Seeds | GB66592 | 2016-057 | Balibisiana | *Musa balbisiana* | Camarines Norte |
| Seeds | GB66592 | 2016-057 H2 | Moko | *Musa balbisiana* | Camarines Norte |
| Seeds | GB66648 | 2017-040 | Balbisiana | *Musa balbisiana* | Quezon |
| Seeds | GB67532 | 2017-042 | Agutay | *M. acuminata*<br>*subsp. errans* | Laguna |
| Seeds | GB67541 | 2017-048 | Pik-iw | *Musa balbisiana* | Quezon |

| Plant Source Type | GB/ PHL Number | Collection Number | Local name | Scientific name | Collection Site |
| --- | --- | --- | --- | --- | --- |
| Seeds | GB67548 | 2017-055 | Balbisiana | *Musa balbisiana* | Quezon |
| Seeds | GB 71332 | 2021-029 | Agutay | *M. acuminata*<br>*subsp. errans* | Laguna |
| Tissue culture |  | 1999-058 | Butuhan | *Musa balbisiana* | Laguna |
| Tissue culture |  | 1999-088 | Butuhan | *Musa balbisiana* | Davao Del Sur |

**Figure 1.**
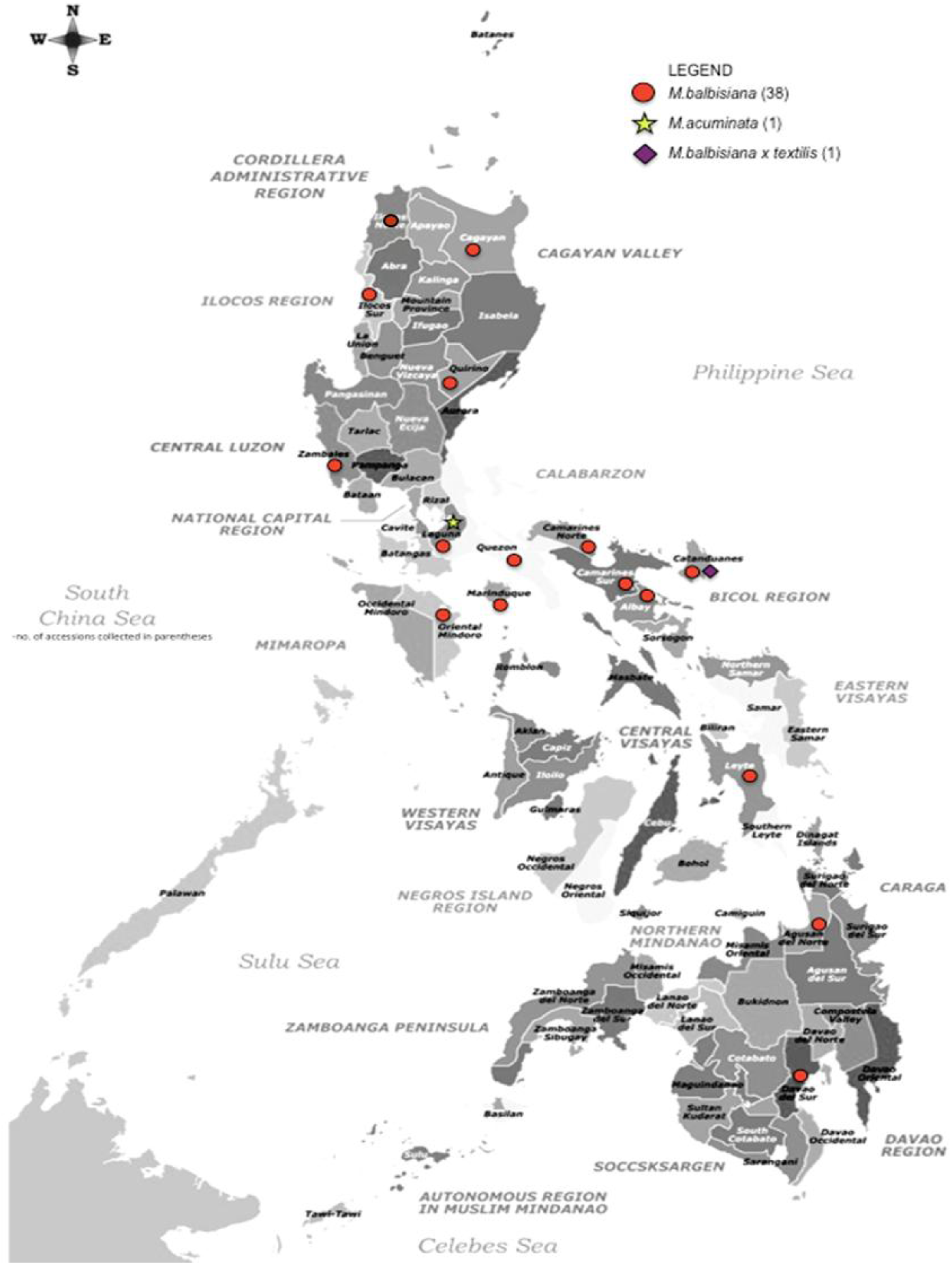
Philippine map showing the places of collection (in red dots) of different *Musa balbisiana* accessions.43

### Collection of BBTV inoculum and aphid vector

Reference BBTV isolate REF2-LAG was obtained from banana cv. Lakatan with typical BBTV symptoms from Los Baños, Laguna, Philippines. An 861 nt fragment of the DNA-R segment of the isolate was obtained using primer pair BBTVREP-F and BBTVREP-R (Islam et al. 2010). The resulting sequence was deposited in GenBank with accession code ON038409. For the banana aphid vector (*Pentalonia nigronervosa*), a colony was derived from a virus-free wingless nymph reared on virus-free banana cv. Lakatan plants (Parna & Agarwala, 2010). Modified Dellaporta minipreparation procedure was used in the extraction of aphid DNA (Dellaporta et al., 1983). PCR Primers LepF and LepR (Hebert et al. 2004) were used to amplify the mitochondrial cytochrome c oxidase I gene. The PCR conditions used are as follows: 94°C for 2 min, 35 cycle of 94°C for 1 min, 56°C for 30 s and 72°C for 1 min, and final extension at 72°C for 5 min. The amplicon was sequenced, and the sequence deposited in GenBank under accession number ON077361. The BBTV isolate and aphid colony were kept in 32-mesh insect-proof screen cages under greenhouse conditions.

### Screening of wild Musa test plants for BBTV resistance

Accessions from seeds had a total of 60 replicates while tissue cultured accessions had 25 replicates. A standard transmission protocol of BBTV was performed using aphids from a single colony and the BBTV reference isolate. Aphid transmission of BBTV followed the protocol of Thomas and Dietzgen (1991) with a few modifications (Dijkstra and Jager, 1998; Anhalt and Almeida, 2008). Wingless aphids were transferred into a sterilized closed container and starved for four hours. Starved aphids were then transferred to a banana cv. Lakatan plant infected with the BBTV reference isolate for an 18 to 24-hour acquisition access period (AAP). Twenty viruliferous aphids were then transferred to each test plant and allowed to feed for a 24 to 30-hour inoculation access period (IAP). Ten banana cv. Lakatan plants (dela Cruz et al., 2008) were used as positive controls per batch of inoculations. Inoculated test plants were sprayed with insecticide to eliminate the aphids after the IAP. Samples were maintained inside an insect-proof or 32 mesh screen cage in greenhouse conditions and regularly observed for symptom appearance.

### Confirmation of resistance of wild banana accessions from greenhouse trial

Wild banana accessions with 0% BBTD infection six months post-inoculation in greenhouse were then transferred to a field with high incidence of BBTD for further evaluation. Basic land preparation was performed such as mowing, plowing, rotavating and furrowing prior to transplanting. Wild *Musa* accessions were transplanted to the field per batch of screening. As a susceptible check, six-month-old virus-free banana cv. Lakatan plants were also planted parallel to test plants with the same number of replicates.

Screened *M. balbisiana* accessions in the field with suckers and producing bunches with viable seeds were subjected to re-screening against BBTV. The corms of the suckers of accessions 2016-018, 1998-087, 2016-013, 2013-155, 2016-051, 2016-003, and 2013-101 were macropropagated in large drums in an insect-proof cage and a total of four replicates per accession were screened against BBTV. On the other hand, seeds from accessions 2016-003, 2013-155, 2013-156, 1998-078, 1997-276 were harvested and grown in seedbeds. Fifteen seedlings per accession served as replicates. Test plants were inoculated with BBTV using the standard transmission protocol and monitored as outlined above. The trial was performed under greenhouse conditions.

### Detecion of BBTV

The presence of BBTV in test plants in the greenhouse was determined by PCR. Detection was conducted at three and six months after inoculation before proceeding to the field trial. Then, for accessions in the field, samples were taken from 10 randomly selected plants per *Musa* accession every six months from transplanting. Leaf samples were taken from the second youngest leaf with portions of the midrib (Thomas et al., 1994). About 0.5g per sample was homogenized in a 100 × 150 mm polyethylene bag with a 3 mL extraction buffer (0.05 M Tris-HCl, 0.5% sodium sulfite, and 2.5% non-fat milk). One ml of leaf extract was transferred into a sterile 1.5 ml tube and centrifuged for 5 min at 10,000 rpm. The supernatant was used as a template for PCR assay targeting the DNA-R of BBTV using BBT1 and BBT2 primer pair (Thomson and Dietzgen 1991). Exactly 2 µl of template was added into a PCR cocktail containing 1x PCR buffer, 1.76 mM MgCl^2^, 0.2 mM dNTPs, 0.2 µM of forward BBT1 (3’ CTCGTCATGTGCAAGGTTATGTCG 5’) and reverse primer BBT2, (5’ GAAGTTCTCCAGCTATTCATCGCC 3’), 1 U Taq DNA polymerase and DEPC water to attain a final volume of 15 µl. Samples were run in a Mycycler thermal cycler (Bio-Rad, USA) using the following PCR conditions: 94°C for 3 min, 35 cycles of 94°C for 30 s, 56°C for 1 min and 72°C for 1 min, and a final extension of 3 min at 72°C. The presence of BBTV was indicated by the amplification of the 349 bp band visualized by gel electrophoresis with ethidium bromide staining. PCR assays were duplicated.

## Results

### Screening of wild *Musa* test plant accessions

The positive control banana cv. Lakatan was the first to show symptoms of BBTD. Marginal chlorosis and dark green streaks along midrib were observed between 15-21 days after inoculation (DAI). Among wild banana accessions, only *M*. *acuminata* subsp. *errans* accessions, 2017-042 and 2021-029, expressed initial symptoms of BBTD at 21-28 DAI as shown in figure 2. By six weeks after inoculation, 100% disease incidence was recorded in *M*.*acuminata subsp*. *errans* showing confirmatory symptoms of BBTD such as dot and dash flecks, narrow and erect emerging leaves with marginal chlorosis and bunchy top. Meanwhile, none of the 34 *M. balbisiana* accessions showed any symptoms of the disease even at six-month post-inoculation (Figure 3). At three months post-inoculation, *M*. *acuminata* subsp. *errans* accessions and cv. Lakatan recorded a 100% BBTV incidence based in PCR assays. All *M*. *balbisiana* entries were negative after repeated PCR assays as indicated in Table 2.

**Table 2.** Result of greenhouse and field screening of wild banana against BBTV

| Collection Number | Scientific name | Source | Greenhouse<br>BBTV % Infection | Field<br>BBTV % Infection | Location in the field |
| --- | --- | --- | --- | --- | --- |
| <u>Batch 1 and 2</u> |  |  |  |  |  |
| 2013-155 | *Musa balbisiana* | Seeds | 0 | 0 | Plot 1 |
| 2016-003 | *Musa balbisiana* | Seeds | 0 | 0 | Plot 1 |
| 2013-159 | *Musa balbisiana* | Seeds | 0 | 0 | Plot 2 |
| 2016-051 | *Musa balbisiana* | Seeds | 0 | 0 | Plot 2 |
| Positive Control<br>Lakatan | *M. acuminata* | Tissue<br>Cultured | 100% | 100% | n/a (100%*) |
| <u>Batch 3</u> |  |  |  |  |  |
| 2013-101 | *Musa balbisiana* | Seeds | 0 | 0 | Plot 3 |
| 2013-156 | *Musa balbisiana* | Seeds | 0 | 0 | Plot 3 |
| 2017-040 | *Musa balbisiana* | Seeds | 0 | 0 | Plot 3 |
| **2017-042** | ***M. acuminata***<br>***subsp. errans*** | **Seeds** | **100%** | **NT\*\*** | **Greenhouse Trial**<br>**only** |
| Positive Control<br>Lakatan | *M. acuminata* | Tissue<br>Cultured | 100% | 100% | n/a (100%*) |
| <u>Batch 4</u> |  |  |  |  |  |
| 1998-085 | *Musa balbisiana* | Seeds | 0 | 0 | Plot 4 |
| 1998-315 | *Musa balbisiana* | Seeds | 0 | 0 | Plot 4 |
| Positive Control<br>Lakatan | *M. acuminata* | Tissue<br>Cultured | 100% | 100% | n/a (100%*) |

\*% Mortality rate \*\*NT- Not tested

Table 2. Result of greenhouse and field screening of wild banana test plants against BBTV (Continuation...)
| Collection Number | Scientific name | Source | Greenhouse<br>BBTV % Infection | Field<br>BBTV % Infection | Location in the field |
| --- | --- | --- | --- | --- | --- |
| <u>Batch 5</u> |  |  |  |  |  |
| 1999-088 | *Musa balbisiana* | Tissue Cultured | 0 | 0 | Plot 5 |
| 1998-058 | *Musa balbisiana* | Tissue Cultured | 0 | 0 | Plot 5 |
| 2010-047 | *Musa balbisiana* | Seeds | 0 | 0 | Plot 5 |
| 2016-013 | *Musa balbisiana* | Seeds | 0 | 0 | Plot 5 |
| 2016-018 | *Musa balbisiana* | Seeds | 0 | 0 | Plot 5 |
| 1998-078 | *Musa balbisiana* | Seeds | 0 | 0 | Plot 5 |
| Positive Control Lakatan | *M. acuminata* | Tissue Cultured | 100% | 100% | n/a (100%*) |
| <u>Batch 6</u> |  |  |  |  |  |
| 2016-052 | *Musa balbisiana* | Seeds | 0 | 0 | Plot 6 |
| 2016-018 | *Musa balbisiana* | Seeds | 0 | 0 | Plot 6 |
| 1997-276 | *Musa balbisiana* | Seeds | 0 | 0 | Plot 6 |
| Positive Control Lakatan | *M. acuminata* | Tissue Cultured | 100% | 100% | n/a (100%*) |
| <u>Batch 7</u> |  |  |  |  |  |
| 2016-048 H1 | *Musa balbisiana* | Seeds | 0 | 0 | Plot 7 |
| 2016-051 H1 | *Musa balbisiana* | Seeds | 0 | 0 | Plot 7 |
| 1998-633 | *Musa balbisiana* | Seeds | 0 | 0 | Plot 7 |
| Positive Control Lakatan | *M. acuminata* | Tissue Cultured | 100% | 100% | n/a (43%*) |

\*% Mortality rate

Table 2. Result of greenhouse and field screening of wild banana test plants against BBTv (Continuation...)
| Collection Number | Scientific name | Source | Greenhouse<br>BBTV % Infection | Field<br>BBTV % Infection | Location in the field |
| --- | --- | --- | --- | --- | --- |
| **Batch 8** |  |  |  |  |  |
| 2013-155 H2 | *Musa balbisiana* | Seeds | 0 | 0 | Plot 8 |
| 2016-056 H3 | *Musa balbisiana* | Seeds | 0 | 0 | Plot 8 |
| 1998-315 H2 | *Musa balbisiana* | Seeds | 0 | 0 | Plot 8 |
| 1998-575 | *Musa balbisiana* | Seeds | 0 | 0 | Plot 8 |
| 2013-100 | *Musa balbisiana* | Seeds | 0 | 0 | Plot 8 |
| 2016-051 H2 | *Musa balbisiana* | Seeds | 0 | 0 | Plot 8 |
| Positive Control Lakatan | *M. acuminata* | Tissue<br>Cultured | 100% | 100% | Plot 8 (50%*) |

\*% Mortality rate

Table 2. Result of greenhouse and field screening of wild banana test plants against BBTv (Continuation...)
| Collection Number | Scientific name | Source | Greenhouse<br>BBTV % Infection | Field<br>BBTV % Infection | Location in the field |
| --- | --- | --- | --- | --- | --- |
| **Batch 9** |  |  |  |  |  |
| 2017-055 | *Musa balbisiana* | Seeds | 0 | 0 | Plot 9 |
| 2013-100 H2 | *Musa balbisiana* | Seeds | 0 | 0 | Plot 9 |
| 2016-057 H2 | *Musa balbisiana* | Seeds | 0 | 0 | Plot 9 |
| 1998-453 | *Musa balbisiana* | Seeds | 0 | 0 | Plot 9 |
| 2016-053 | *Musa balbisiana* | Seeds | 0 | 0 | Plot 9 |
| 2017-048 | *Musa balbisiana* | Seeds | 0 | 0 | Plot 9 |
| 1998-456 | *Musa balbisiana* | Seeds | 0 | 0 | Plot 9 |
| 2021-029 | *M. acuminata*<br>*subsp. errans* | Seeds | 100% | NT** | Greenhouse Trial<br>only |
| Positive Control Lakatan | *M. acuminata* | Tissue<br>Cultured | 100% | 100% | Plot 9 |

\*% Mortality rate \*\*NT- Not tested

**Figure 2.**
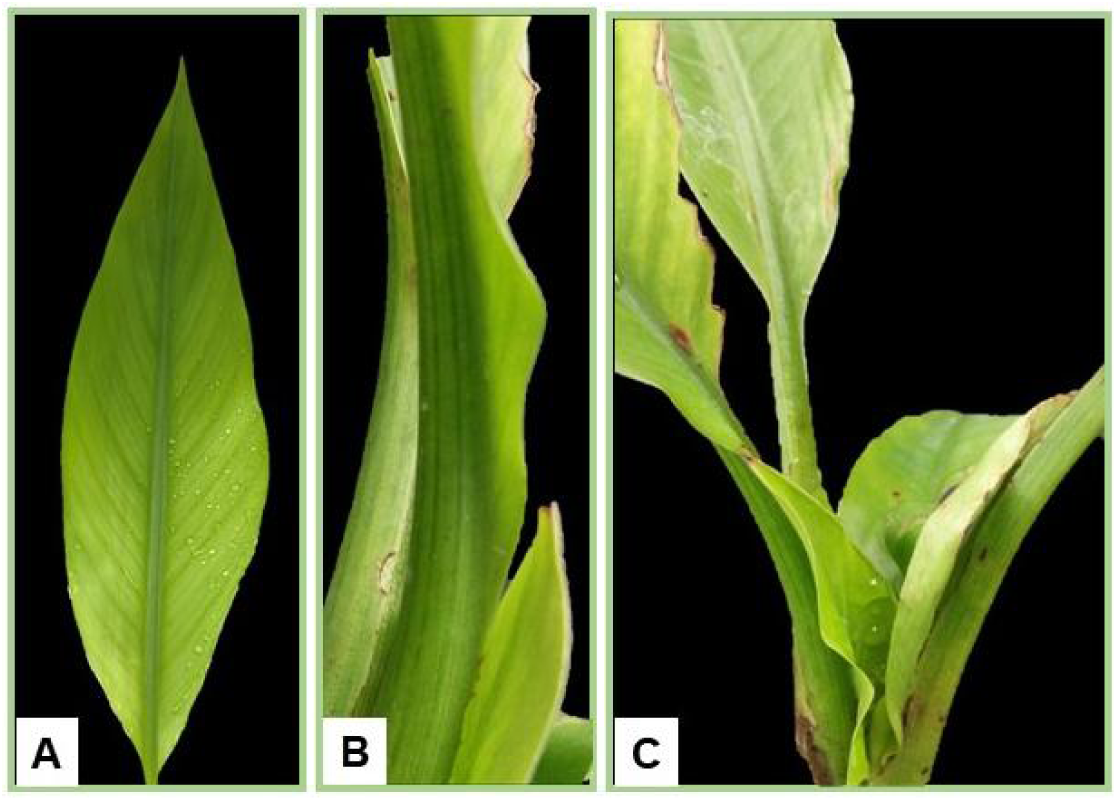
BBTV symptoms on ‘Agutay’ (*M. acuminata* ssp. *errans*): (a) chlorosis on young leaves, (b) dot and dash flecks from midrib to petiole (c) bunchy top

**Figure 3.**
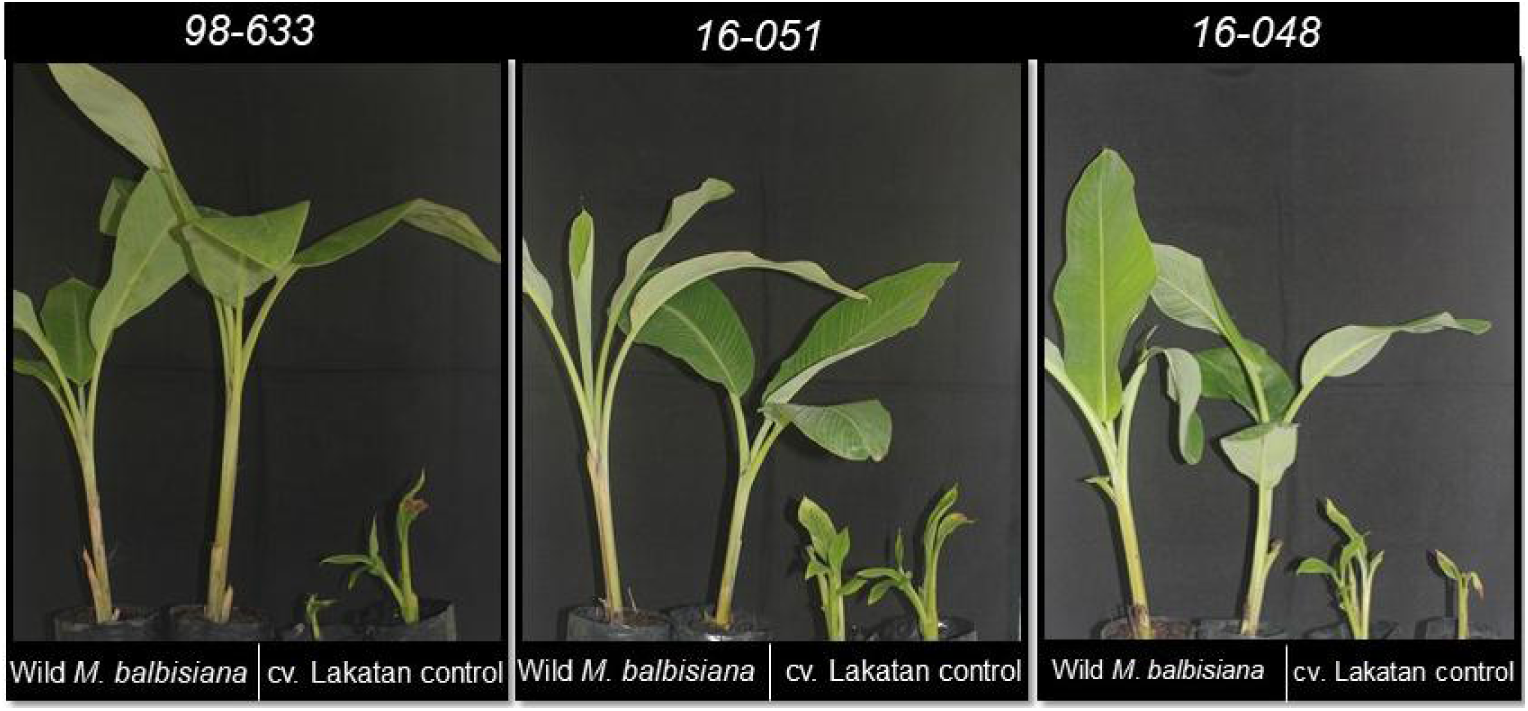
Asymptomatic *M*. *balbisiana* (MB) accessions 1998-633 (a), 2016-051 (b), and 2016-048 (c) in comparison with the susceptible control banana cv. Lakatan (right).

### Confirmation of resistance of wild M. balbisiana accessions (Field Trial)

All 34 Wild *M. balbisiana* accessions from seeds and tissue culture proceeded to the field trial. Healthy tissue-cultured banana cv. Lakatan plants were also planted along the rows of *M. balbisiana* accessions. As early as one month after transplanting, initial symptoms such as j-hook, chlorosis and dark green streaks along the midrib started to appear in cv. Lakatan plants. Symptoms of BBTD became severe with stunting, leaf narrowing, and bunchy top exhibited. Incidence of BBTV in control cv. Lakatan plants per batch of inoculation were indicated in table 2. *M. balbisiana* test plants did not show any symptoms and continued to develop and produce suckers. Vigorous growth was noted even under high disease pressure as shown in in figure 4. Macropropagated plants (figure 5) and seedlings derived from field trial plants were also symptomless.

**Figure 4.**
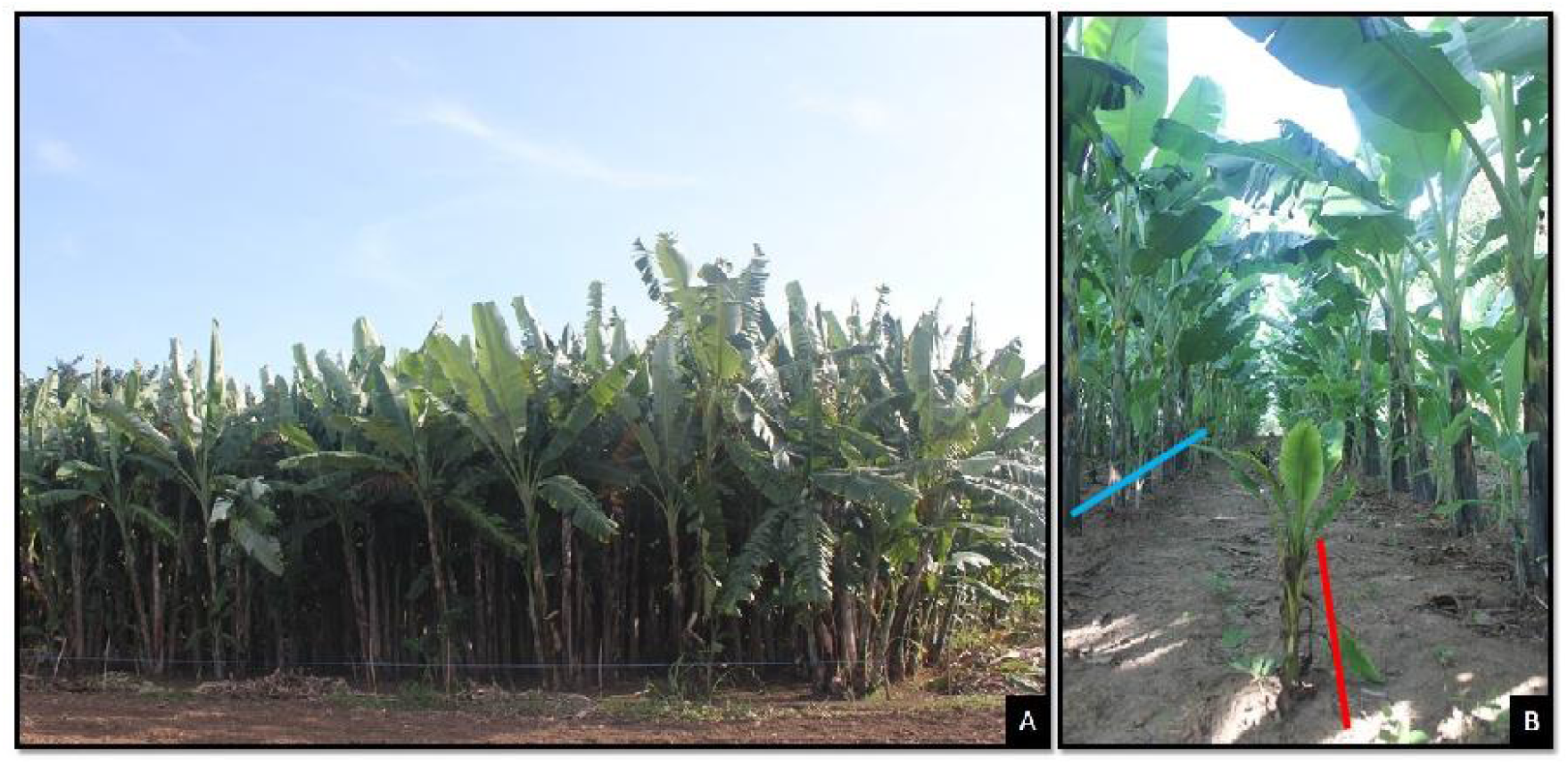
*Musa balbisiana* accessions screening under field trial: (A) oldest standing plot (B) field screening set-up with severely infected cv. Lakatan control. Red line represents susceptible control banana cv. *bla*. Lakatan row. Blue line represents*M. biana* row.

**Figure 5.**
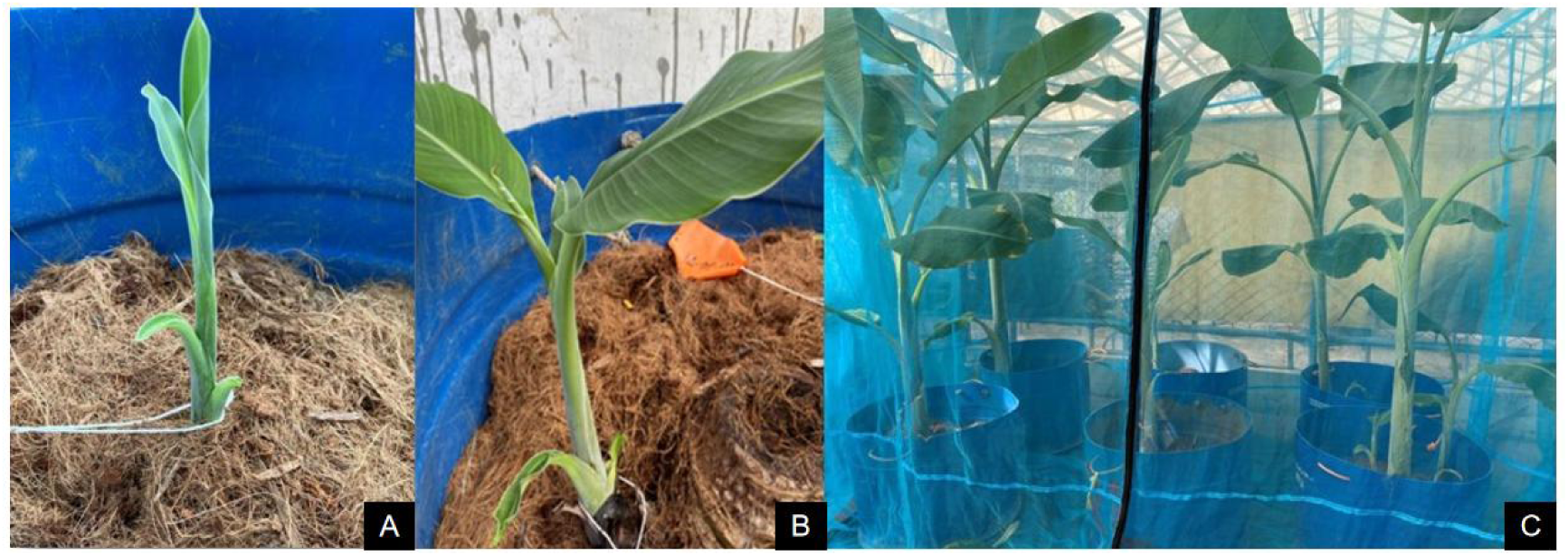
Post screening result: (a-b) Suckers emerging from corms of wild M. balbisiana; (c)suckers developed and remained negative against BBTV

BBTV was not detected by PCR in any replicates of all *M*. *balbisiana* accessions from seeds (n=60) and tissue culture (n=25) that were collected from the field trial while all samples of cv. Lakatan test plants were positive for BBTV infection with 100% incidence at 3 months after inoculation based in PCR. Macropropagated suckers and seedling derivatives of accessions from field trial plants also tested negative for BBTV by PCR.

## Discussion

BBTV causes the most destructive virus disease of bananas. The disease virtually eliminated smallholdings of banana cv. Lakatan in the Luzon region of the Philippines (Molina et al. 2009) and thus most banana produce of the Philippines now comes from the Mindanao region. The most common practice in controlling BBTV in Mindanao is with the use of clean, tissue-cultured plants (Molina et al. 2009). However, the use of virus-free planting materials alone does not guarantee total protection against BBTD, and on-going inspection and eradication programs are challenging to enact and expensive to run. An efficient and sustainable strategy for controlling virus diseases involves the use of resistant varieties. However, there are still no reported varieties of banana resistant to BBTD (Hooks et al. 2009). In this study, the potential of wild *Musa* spp. as natural sources of resistance against BBTV was assessed.

Although *M*. *acuminata* subsp. *errans* accessions were susceptible to BBTV infection with a 100% infection rate and the control banana cv. ‘Lakatan’ showed strong BBTD symptoms and a 100% incidence of BBTV by PCR, all 34 *M. balbisiana* accessions showed no symptoms under greenhouse and field conditions. The virus was also not detected in *M*. *balbisiana* accessions by PCR after multiple rounds of screening. This study has revealed that wild *M. balbisiana* accessions from the Philippines are likely sources of resistance to BBTV. The lack of infection of the *M*.*balbisiana* accessions is unlikely to be due to lack of aphid feeding, as aphid survival numbers on the inoculated plants was high, and indeed some of these *M*. *balbisiana* accessions were used as maintenance hosts for the non-viruliferous aphid colonies.

Susceptibility of *Musa* cultivars has often been found to be related to genomic grouping, with cultivars containing a B genome component derived from *M*. *balbisiana* (BB) often showing a level of tolerance or resistance to BBTV (Ngatat et al., 2017; Jose; 1981, Muharam, 1984; Espino et al., 1993; Niyongere et al., 2011; Sachter-Smith, 2015; Hapsari and Masrum, 2012; Latifah et al., 2021). This association is not always consistent, however, and some A genome cultivars such as Gros Michel (AAA) show resistance to infection by BBTV while B genome-containing cultivars can vary markedly in their reaction (Niyongere et al., 2011; Ngatat et al., 2017; Magee, 1948). Thus, there may be several sources of resistance or tolerance to BBTV in *Musa* germplasm. The results presented here support the hypothesis that a source of resistance to BBTV is present in the B genome of wild *M*.*balbisiana* and that it has been transferred to many edible cultivars that contain a proportion of B genome. However, chromosome structural changes including translocations are known within wild banana species and in edible interspecific hybrids (Šimoníková et al 2022), so it is likely that resistance genes present on a chromosome in one cultivar may not be present in the same chromosome of another. The source of resistance describe in this work appears to be widely distributed in *M*. *balbisiana* from the Philippines.

The centre of origin for *M*.*balbisiana* is thought to range from north-east India, the northern limits of south-east Asia, southern China, to the Philippines (Perrier et al, 2009), though presence in the Philippines may be as feral populations introduced by humans (De Langhe et al 2015). Interestingly the centre of origin for the BBTV pathosystem where the maximum degree of BBTV diversity is found (Rao et al., 2017; Stainton et al., 2015) also coincides with this region. One would expect that resistance to BBTV may have also evolved here.

Although the BBTV-resistant wild *M*. *balbisiana* from the Philippines are seeded, and thus not directly suitable as an edible banana, they have alternative uses. Filipinos prefer the flower bud of wild *M*. *balbisiana* rather than cultivated bananas in cooking. Its thick and green leaves are also marketed in the region for culinary purposes.

Breeding of banana continues with the goal of incorporating resistance to diseases, abiotic stresses and other desirable morphological traits (Wilson et al. 2020). A widely used scheme in banana breeding today is the crossing of triploid and diploid banana (3×/2×) followed by tetraploid and diploid or 4×/2× (Jenny 2002; Vezina 2014). *M*. *balbisiana* accessions can be used as parents for conventional breeding and hand pollinated bunches can produce over 40,000 seeds per bunch. *M. balbisiana* is recognized as a source of useful traits within *Musa* including vigorous growth, strong suckering, drought tolerance and resistance to abiotic stresses (Bakry et al 2021).

With the use of current molecular tools, investigation on molecular mechanisms of resistance in wild *M*. *balbisana* accessions can be conducted (Solomon-Blackburn & Barker 2001; Sharma et al. 2018). The findings of the present study have led to the implementation of a local research project on fast-tracking the development of BBTV-resistant banana cultivars using modern biotechnology tools. A next-generation sequencing strategy (RNA-seq) is currently being performed to find alleles related to BBTD resistance. Discoveries will be further applied in assisting DNA marker development to aid the ultimate development of genetically modified resistant bananas.

## Conclusion

Thirty four wild *M. balbisiana* (BBw) accessions exhibited total resistance to BBTV as shown by the absence of symptoms and negative molecular detection of BBTV under greenhouse and field evaluation. The study confirms that resistance against BBTD exists in the wild ecosystem and provides a solid base for future efforts to produce BBTV-resistant banana cultivars. These accessions could be used as a source of resistance for conventional breeding of BBTV-resistant hybrids. The use of molecular tools to investigate the genetic aspect of resistance in these accessions could lead to applications such as marker assisted breeding and incorporation of resistance genes through transgenic approaches.

## Acknowledgments

This work was supported by the Bill & Melinda Gates Foundation [Grant Number INV-010652]. Under the grant conditions of the Foundation, a Creative Commons Attribution 4.0 Generic License has already been assigned to the Author Accepted Manuscript version that might arise from this submission. We thank Mr. Rodelio Pia, Mr. Valentino L. Garcia, Mr. Joey A. Ancheta, Ms. Rizalina L. Tiongco, Ms. Vangelene T. Linga, Ms. Elaine Anne L. Elmido and Mr. Jay-Vee S. Mendoza from Plant Pathology Laboratory and Mr.Jerome Lopez, Ms. Maria B. Adisaz and Ms. Jovy Marie Cezar from the National Plant Genetic Resources Laboratory of the Institute of Plant Breeding, College of Agriculture and Food Science, University of the Philippines Los Baños for their technical assistance and moral support throughout the study.

## Conflict of Interests

The authors declare that they have no conflict of interests.

## Ethical Approval

This article does not contain any studies with human and animal participations performed by the authors

## Notes

### Competing Interest Statement

The authors have declared no competing interest.

